# A test for microbiome-mediated rescue via host phenotypic plasticity in *Daphnia*

**DOI:** 10.1101/2024.08.14.607994

**Authors:** René S. Shahmohamadloo, Amir R. Gabidulin, Ellie R. Andrews, John M. Fryxell, Seth M. Rudman

**Author notes:** **Corresponding authors:** René S. Shahmohamadloo, School of Biological Sciences, Washington State University, 14204 NE Salmon Creek Ave, Vancouver, WA 98686, USA., Seth M. Rudman, School of Biological Sciences, Washington State University, 14204 NE Salmon Creek Ave, Vancouver, WA 98686, USA. **Statement of authorship:** S.M.R. and R.S.S. designed the research. J.M.F. and S.M.R. provided material resources. R.S.S. and S.M.R. conducted the experiment. R.S.S. and E.R.A. completed extractions. A.R.G. and R.S.S. analyzed the data. R.S.S., S.M.R., A.R.G., and E.R.A. wrote the paper. All authors read, amended, and approved the final manuscript. **Data accessibility statement:** The data supporting the results will be archived in the public repository Dryad (DOI will be released upon publication). **Competing interest statement:** The authors declare no competing interests.

## Abstract

Phenotypic plasticity is a primary mechanism by which organismal phenotypes shift in response to the environment. Host-associated microbiomes often exhibit considerable shifts in response to environmental variation and these shifts could facilitate host phenotypic plasticity, adaptation, or rescue populations from extinction. However, it is unclear how much shifts in microbiome composition contribute to host phenotypic plasticity, limiting our knowledge of the underlying mechanisms of plasticity and, ultimately, the fate of populations inhabiting changing environments. In this study, we examined phenotypic responses and microbiome composition in 20 genetically distinct *Daphnia magna* clones exposed to non-toxic and toxic diets containing *Microcystis*, a cosmopolitan cyanobacteria and common stressor for *Daphnia. Daphnia* exhibited significant plasticity in survival, reproduction, and population growth rates in response to *Microcystis* exposure. However, the effects of *Microcystis* exposure on the *Daphnia* microbiome were limited, with the primary effect being differences in abundance observed across five bacterial families. Moreover, there was no significant correlation between the magnitude of microbiome shifts and host phenotypic plasticity. Our results suggest that microbiome composition played a negligible role in driving host phenotypic plasticity or microbiome-mediated rescue.

**One sentence summary:** *Daphnia* exhibits considerable plasticity in individual and population-level responses to a cosmopolitan stressor, yet shifts in microbiome composition are not correlated with the magnitude of this plasticity.

## Introduction

Projected biodiversity losses from anthropogenic global change this century (Urban et al., 2016) necessitate an understanding of the magnitude and mechanisms of adaptive phenotypic responses to environmental change (Bellard et al., 2012; Lavergne et al., 2010). Although organisms can rapidly evolve in response to global change (Hendry & Kinnison, 1999; Hoffmann & Sgrò, 2011; Rudman et al., 2022), environmentally induced phenotypic plasticity—defined as the capacity for a given genotype to express different phenotypes when faced with a different environmental challenge (West-Eberhard, 2003)—is a primary mechanism by which organisms respond to environmental changes (Ghalambor et al., 2007; Merilä & Hendry, 2014). This plasticity can be adaptive, meaning it can enhance an organism’s fitness in response to environmental change, and such adaptive plasticity is a key component of reducing phenotype-environment mismatches that can cause extinction (Bell & Collins, 2008; Bell & Gonzalez, 2009). Yet, the mechanisms that produce adaptive phenotypic plasticity are not always well understood.

The microbiome impacts nearly all aspects of host phenotype (Akbar et al., 2022; Berg et al., 2020; Flandroy et al., 2018; Moran & Baumann, 2000) and extends the host’s genetic repertoire (Decaestecker et al., 2024). Host-associated microbiomes both show sensitivity to environmental changes and have demonstrated diverse impacts on host fitness (Decaestecker et al., 2024). The composition of associated microbial communities can influence host performance and relative fitness (Gould et al., 2018), and intraspecific variation in microbiome composition has been observed to affect host physiology and performance in various taxa (Bolnick et al., 2014; Rudman et al., 2019; J. Wang et al., 2015). This variation and its impact on host phenotypes have spurred considerable speculation about the microbiome’s crucial role in host population persistence(White et al., 2023) and in extending host adaptive capacities (Henry et al., 2021).

Microbiome-mediated plasticity—defined as changes in microbiome composition or function that influence host phenotype—has been proposed as a general mechanism that facilitates evolutionary adaptation by reducing mismatches between host phenotype and the environment (Henry et al., 2021; Kolodny & Schulenburg, 2020). Experimental work on sea anemones suggest that microbiome-mediated plasticity can enhance thermal acclimation and that the benefits of such acclimation can be transmitted to the next generation, suggesting microbiota as a mechanism of rapid adaptation to environmental changes (Baldassarre et al., 2022). Coral microbiomes too may shift to protect against extreme challenges like heat stress and pathogens, underpinning the health and resilience of reef ecosystems (Bourne et al., 2016; Webster & Reusch, 2017). Research using *Caenorhabditis elegans* in a novel environment—though with limited population-level replication—also suggests that adaptation to new conditions can be jointly influenced by host genetics and microbiome composition, highlighting the complex interactions between host and microbiome in shaping evolutionary trajectories (Petersen et al., 2023).

Cases where microbiome-mediated plasticity impact host population growth and persistence under challenging environments have received particular attention. This has led to the concept of microbiome rescue—defined here as changes in microbiome abundance, composition, or activity that improves host fitness and decreases the likelihood of extinction (Mueller et al., 2020)—and speculation that it may be an important process for the maintenance of biodiversity in rapidly changing environments (Mueller et al., 2020; Shade, 2023). Microbiomes can play a critical role in helping hosts cope with various environmental challenges, including toxicants, by potentially modulating detoxification mechanisms and stress responses (Adamovsky et al., 2018; Kikuchi et al., 2012; G.-H. Wang et al., 2020). However, while these studies highlight the potential for microbiomes to influence host responses to stress, they do not provide direct evidence of microbiome rescue in populations. Limited empirical evidence remains linking shifts in microbiome composition to significant impacts on host population dynamics outside of obligate symbiosis, leaving considerable uncertainty about the strength of any such beneficial effects or the potential for rescue (Mason, 2020).

Empirical tests of the contributions of shifts in the microbiome to adaptive plasticity are needed to determine whether microbiomes facilitate population persistence or rescue in response to environmental stress. *Daphnia* (water fleas) exhibit observable and quantifiable phenotypic changes in response to environmental fluctuations that are crucial to population dynamics, including alterations in survival rates, morphology, and reproductive strategies (Boersma et al., 1998). The interaction between *Daphnia* and harmful algal blooms (HABs) of the cyanobacterium *Microcystis*—a global environmental challenge (Harke et al., 2016)—is both important for host fitness and environmental health (Hairston et al., 2001; Isanta-Navarro et al., 2021; Shahmohamadloo, Poirier, et al., 2020; Shahmohamadloo, Simmons, et al., 2020). Microbiome transplant experiments have demonstrated that both host genotype and gut microbiota can mediate tolerance in *Daphnia* to *Microcystis* (Macke et al., 2017); host genotypic variation in tolerance disappears when *Daphnia* are made germ-free and inoculated with a standardized microbiome (Macke et al., 2017). In another reciprocal transplant experiment, *Daphnia* performed better when receiving a microbiome from their source region when exposed to toxic *Microcystis*, indicating microbiome-mediated local adaptation in stress tolerance (Houwenhuyse et al., 2021). This effect was most pronounced when donor microbiomes were pre-exposed to toxic cyanobacteria, and it also depended on the pond and genotype of the *Daphnia* (Houwenhuyse et al., 2021). More broadly, the *Daphnia* microbiome exhibits considerable plasticity in both diversity and composition, influenced by the environmental bacterial community and host genotype, among other potential environmental drivers (Callens et al., 2020; Hegg et al., 2021; Houwenhuyse et al., 2021).

Prior work on *Daphnia*-*Microcystis* interactions has been instrumental in revealing the mechanisms by which microbiomes can impact hosts, but understanding the implications of these interactions for host plasticity and population dynamics requires different approaches. First, determining the extent of *Daphnia* microbiome shifts, and whether these shifts are repeated and deterministic, is critical to understanding how robust the effects of microbiome changes may be on host phenotypes (Härer & Rennison, 2022; Stuart et al., 2017). Second, explicitly testing whether the magnitude or direction of microbiome shifts are correlated with overall phenotypic plasticity is key to understanding the contribution of microbiome change to host phenotypic response (Kolodny & Schulenburg, 2020). Finally, measuring the magnitude of an observed shift on host fitness due specifically to microbiome alteration is critical in evaluating the microbiome rescue hypothesis (Mueller et al., 2020; Shade, 2023). Doing so requires measurements of the relationship between microbiome shifts and parameters central to population dynamics. Together these data can provide a test of the role of microbiome shifts in host phenotypic plasticity and population dynamics in changing environments.

To address key questions regarding the role of the microbiome in facilitating host plasticity and rescue, we conducted an algal toxicity and microbiome study on 20 genetically distinct clones of *Daphnia magna* collected from a single lake. We first catalog the impacts of *Microcystis* exposure on *Daphnia* phenotypes and population growth rate for each genotype. We then tested the following questions: 1) What is the magnitude of *Daphnia* phenotypic response to *Microcystis* exposure? 2) Does *Microcystis* exposure alter *Daphnia* microbiome composition and abundance and, if so, are these changes parallel and deterministic across different genotypes, under varying treatments? 3) Is the magnitude and/or direction of shifts in the microbiome in response to *Microcystis* correlated with the magnitude of adaptive phenotypic plasticity in *Daphnia*? To answer these questions, we conducted a 21-d chronic toxicity exposure experiment across two common gardens (non-toxic, *Chlorella*-only; and toxic, 3:1 ratio of *Chlorella* to *Microcystis*) using 20 *D. magna* clones. Clonal replication allows for an assessment of the effects of microbiome shifts across multiple genetic backgrounds and enables projections of population-level responses. We use these fitness-associated phenotypes in each of 20 clones to parameterize population projection models and test for the effects of shifts in the microbiome on host plasticity.

## Methods

### Daphnia magna field collection and culturing

In late spring and early summer, 20 genotypes of *D. magna* were obtained from ‘Langerodevijver’ (LRV; 50° 49’ 42.08’’, 04° 38’ 20.60’’), a small lake situated within the nature reserve of Doode Bemde, Vlaams-Brabant, Belgium (Orsini et al., 2012). LRV, with a surface area of 140,000 m^2^ and a maximum depth of 1 m, features a single basin and experiences seasonal HABs of *Microcystis*, a common occurrence in lakes worldwide. Additionally, LRV harbors a large population of *D. magna*. Parthenogenetic lines of each genotype were maintained for over five years (approximately 300 generations) in continuous cultures at 20 ºC, utilizing UV-filtered dechlorinated municipal tap water enriched with 2 mg C L^-1^ of the green alga *C. vulgaris* (strain CPCC 90; Canadian Phycological Culture Centre, Waterloo, ON, Canada). Culturing of *C. vulgaris* was carried out using COMBO medium (Kilham et al., 1998). Filters with a pore size of 0.22 µm were placed at both the input and output of the aeration system to prevent any bacterial contamination.

### Microcystis aeruginosa culturing

In accordance with our previously outlined methodology (Shahmohamadloo et al., 2019), *M. aeruginosa* (strain CPCC 300; Canadian Phycological Culture Centre, Waterloo, ON, Canada) was cultivated in BG-11 media and maintained in a growth chamber under sterile conditions at a constant temperature of 21 ± 1 ºC, illuminated with cool-white fluorescent light at an intensity of 600 ± 15 lx, and subjected to a photoperiod of 16:8 h light:dark. The culture was allowed to grow undisturbed for at least one month before being prepared for the 21-d chronic study. *M. aeruginosa* CPCC 300 is known to produce microcystin-LR (CAS: 101043-37-2, C_49_H_74_N_10_O_12_) and its desmethylated form [D-Asp^3^]-microcystin-LR (CAS: 120011-66-7, C_48_H_72_N_10_O_12_), which are prevalent in freshwater ecosystems and exhibit toxicity to zooplankton (Chorus & Welker, 2021; Harke et al., 2016).

To facilitate testing on *D. magna*, an aliquot of the stock was inoculated into 100% COMBO medium two weeks prior to test initiation, where it was cultured until reaching a cell density of 1.25 ± 0.02 × 10^7^ cells mL^-1^. This medium was chosen because it supports the growth of both algae and cyanobacteria while remaining non-hazardous to zooplankton (Kilham et al., 1998). Filters with a pore size of 0.22 µm were placed at both the input and output of the aeration system to prevent any bacterial contamination.

### Gut microbiome experiment

We evaluated shifts in the microbiome to *M. aeruginosa* using 20 genotypes of *D. magna*. Phenotypic responses measured include survival, reproduction (number of offspring produced), and the timing and number of broods.

In preparation for this investigation, we individually housed one adult female *D. magna* per genotype in separate 50-mL glass tubes containing COMBO medium and *C. vulgaris* at a concentration of 2 mg C L-1. Daily monitoring was conducted to observe reproduction. Neonates of *D. magna* born within 24 h were gathered from each genotype and individually placed into 50-mL glass tubes, following the previously described procedure. This process resulted in 10 replicates per genotype and a total of 200 tubes. These 200 *D. magna*, representing 20 genotypes, served as the founding mothers for this study. All *D. magna* were maintained under constant environmental conditions, including a temperature of 21 ± 1 ºC, cool-white fluorescent light at an intensity of 600 ± 15 lx, and a photoperiod of 16:8 h light:dark.

A 21-day chronic toxicity study was performed following previously described methods (Houwenhuyse et al., 2021; OECD, 2012). *D. magna* neonates from each of the 20 genotypes, born within a 24-hour period, were collected. Groups of 15 neonates were placed in 1-L glass jars containing 750 mL of UV-filtered water, with triplicates for each genotype. Each jar was then assigned to its respective feeding treatment. Two common gardens were included: *Chlorella*-only (non-toxic diet) and 3:1 *Chlorella*:*Microcystis* (toxic diet). Following these ratios, all *D. magna* were fed 2 mg C L^-1^, corresponding to 3 × 10^6^cells total, consistent with previous studies exposing daphnids to dietary combinations of green algae and cyanobacteria (Isanta-Navarro et al., 2021; Rohrlack et al., 2005; Shahmohamadloo, Poirier, et al., 2020). In total, 1,800 animals were used for this study (i.e., 3 replicates per genotype, 900 animals per treatment).

As this was a semi-static test, solutions were refreshed 3 × per week on Mondays, Wednesdays, and Fridays. This process involved transferring *D. magna* from the old to the new glass jar and providing *D. magna* with a food supply consisting of 3 × 10^6^ cells (i.e., the 3:1 treatment received 2 × 10^6^ *C. vulgaris* cells and 1 × 10^6^ *M. aeruginosa* cells, corresponding to 2 mg C L^-1^). Throughout this period, daily survival rates, reproductive output, and the timing and number of broods were recorded to evaluate potential interactions between genotype and treatment effects. The study was conducted under 400– 800 lx cool-white fluorescent light at a temperature of 20 ± 1 °C with a 16:8 light:dark cycle.

At 21 days, *D. magna* replicates were transferred to sterile-filtered tap water for 24 h to eliminate food particles from the gut and environmental bacteria from the carapace and filter apparatus (Houwenhuyse et al., 2021). Subsequently, *D. magna* guts were extracted using dissection needles under a stereomicroscope and transferred into an Eppendorf tube filled with 10 µL sterile milliQ water, after which they were stored in −80 °C for future DNA extraction.

### Library preparation and sequencing

To analyze the gut microbial communities of *D. magna* at the conclusion of the 21-d experiment, DNA extractions were performed using the NucleoSpin Soil DNA kit (Macherey-Nagel, Düren, Germany). Prior to bead beating samples according to kit instructions, each *D. magna* gut was pressed using a stainless steel probe. DNA was eluted using 50 µL of 5 mM Tris/HCl at pH 8.5. Subsequent DNA quantification was conducted with 5 µL per sample of eluted DNA on a Qubit (Invitrogen, Massachusetts, United States) using a dsDNA broad-range assay. Samples that had low DNA concentrations were vacufuged before sequencing to increase concentration. Isolated DNA was quantified (Equalbit 1x dsDNA HS Assay kit) and amplified using primers covering the V3-V4 hypervariable 16s rRNA region. Library quality was assessed and libraries were dual indexed before Equimolar pooling based on QC values. Pooled libraries were sequenced on an Illumina MiSeq with a 250 bp read length configuration to a depth of 0.3M reads for each sample.

Data were imported using demux single end forward reads with initialization of primer trimming followed by high-resolution DADA2 (Callahan et al., 2016) filtering at truncation quality 20 (if nucleotide of quality 20 is detected, the sequence from then on is truncated). Reads were then filtered using default QIIME2 parameters (Bolyen et al., 2018). We retained an average of 16941 reads per sample (Quantile range: 1% = 7237, 25% = 14943, 50% = 17006, 75% = 19835, 100% = 25629).

Taxonomy was assigned using greenegene full length 16s rRNA backbone database with a scikit-learn naive Bayes machine-learning classifier (Bokulich et al., 2018). Before transitioning into the *phyloseq* package in R (R Core Team, 2022), we generated an unrooted phylogenetic tree to improve the accuracy of the downstream analysis. In *phyloseq*, the combined dataset was first filtered by taxa that were represented with less than 5 total taxonomy classifications in the whole sample set, then rarefied at a depth of 20,000 ASV. Data was aggregated to bacterial class for analyses of relative abundance and differential abundance analysis graphs (Figure 3).

### Water sampling for analysis

Water samples were collected from each treatment at the beginning of the test, during solution changes, and at the conclusion to measure cell concentrations and standard water parameters. These samples were quick-frozen at -80 °C and subsequently analyzed for cyanobacterial toxins, which have been reported previously (Shahmohamadloo et al., 2023). Briefly, water parameters recorded before and after solution renewals remained within the specified test criteria. No indications of hypoxia-induced stress on *D. magna* behavior were observed. Mortality and immobilization were initially detected within 48 hours after exposure to the toxic treatments and persisted throughout the 21-d test period. This outcome was anticipated, given that the selected toxic concentrations can range from sublethal (Shahmohamadloo, Poirier, et al., 2020) to lethal (Ferrão-Filho et al., 2000; Lürling & van der Grinten, 2003) in *Daphnia* laboratory assays, and are commonly encountered in freshwater ecosystems affected by HABs (Chorus & Welker, 2021).

### Statistical analysis

All analyses were completed in R version 4.2.2 (R Core Team, 2022). To test the magnitude of clonal variation across treatments, we used a Linear Mixed Effects (LME) model for each *D. magna* phenotypic response with ‘clone’ treated as a random effect and ‘treatment’ treated as a fixed effect. We constructed Leslie matrices for each clone in the toxic and non-toxic exposure treatments. The population growth rate (λ) and exponential rate of increase (*r*) based on the Euler-Lotka equation were additionally calculated to estimate the reproductive output of *D. magna* that were exposed to toxic and non-toxic diets. These calculations were constructed using the full 21-d data on *D. magna*. The difference in neonate production between toxic and non-toxic diets were calculated for each ‘clone’ and ‘treatment’ combination and plotted.

Downstream microbiome analyses were conducted in the *phyloseq* package in R. We calculated Shannon diversity index and tested for the effects of algal treatment and clone on α-diversity using the *lme4* package in R. Bray-Curtis and Weighted Unifrac ß-diversity metrics were used to assess the multivariate effects of algal treatment and clone in a permutational multivariate analysis (PERMANOVA) adonis2 function from the *vegan* package in R. ß-diversity distance matrices were treated as response variables with ‘treatment’ + ‘clone’ treated as fixed effects and the number of sampled guts in each replicate treated as a random effect. To identify bacterial differences on the family level between samples associated with algal treatment, we used a differential abundance analysis from the *DESeq2* package in R, where ASVs representing less than 1% of the reads were discarded and p-values were adjusted using Benjamini-Hochberg procedure (Benjamini & Hochberg, 1995). To support our findings from *DESeq2*, we additionally employed the *phylofactor* package in R as an alternative which utilizes phylogenetic data to assess the differences in microbial community composition (Washburne et al., 2017).

We tested for relationships between shifts in the gut microbiome composition and host plasticity across algal treatments. We used the *multivarvector* package in R (Härer & Rennison, 2022) to generate multivariate vectors and angles by connecting the population means of Principal Component Analysis (PCA) scores with clones across different algal treatments (20 total vectors and angles) as an empirical measure of microbiome shifts in response to *Microcystis* exposure. The relationship between the magnitude of plasticity in microbial composition and host performance, *r* and delta mean neonate production, was assessed using the goodness of fit of R^2^ extracted from a linear model.

To assess the degree of parallelism in the microbiome shift associated with algal treatment, we used *multivarvector* package, generating the angular value of the parallelism of replicates given toxic and non-toxic diets to then assess the parallel, non-parallel, or anti-parallel relationships of our data. In addition to *multivarvector*, we implemented betadisper from *vegan* to assess the multivariate homogeneity of groups dispersions on our ß-diversity metrics.

## Results

### Daphnia magna plasticity in life-history traits and population projection

Exposure to the toxic diet (3:1 Chlorella:Microcystis) had widespread plastic effects on *Daphnia*, including a significant decrease in survival (F_1,19_ = 10.81, p = 0.0039; Figure 1a). *Daphnia* from the non-toxic diet had a 81.5 ± 1.3% survival rate compared to *Daphnia* from the toxic diet who had a 68.13 ± 3.2% survival rate to the cessation of the study at 21 days. Similarly, we observed a significant effect on neonate production per *Daphnia* (F_1,19_ = 19.56, p = 0.00029; Figure 1b); *Daphnia* reared on non-toxic diets produced 4.76 ± 0.29 neonates per *Daphnia* compared to 3.02 ± 0.23 neonates per *Daphnia* for those reared on toxic diets. Lastly, we observed a significant delay in time to first brood (F_1,19_ = 6.66, p = 0.018; Figure 1c); *Daphnia* reared on non-toxic diets reproduced at 9.52 ± 0.15 days compared to 10.53 ± 0.26 days for those reared on toxic diets. We did not observe significant effects on the total number of broods per *Daphnia* (F_1,19_ = 0.25, p = 0.62) by 21 days.

**Figure 1.**
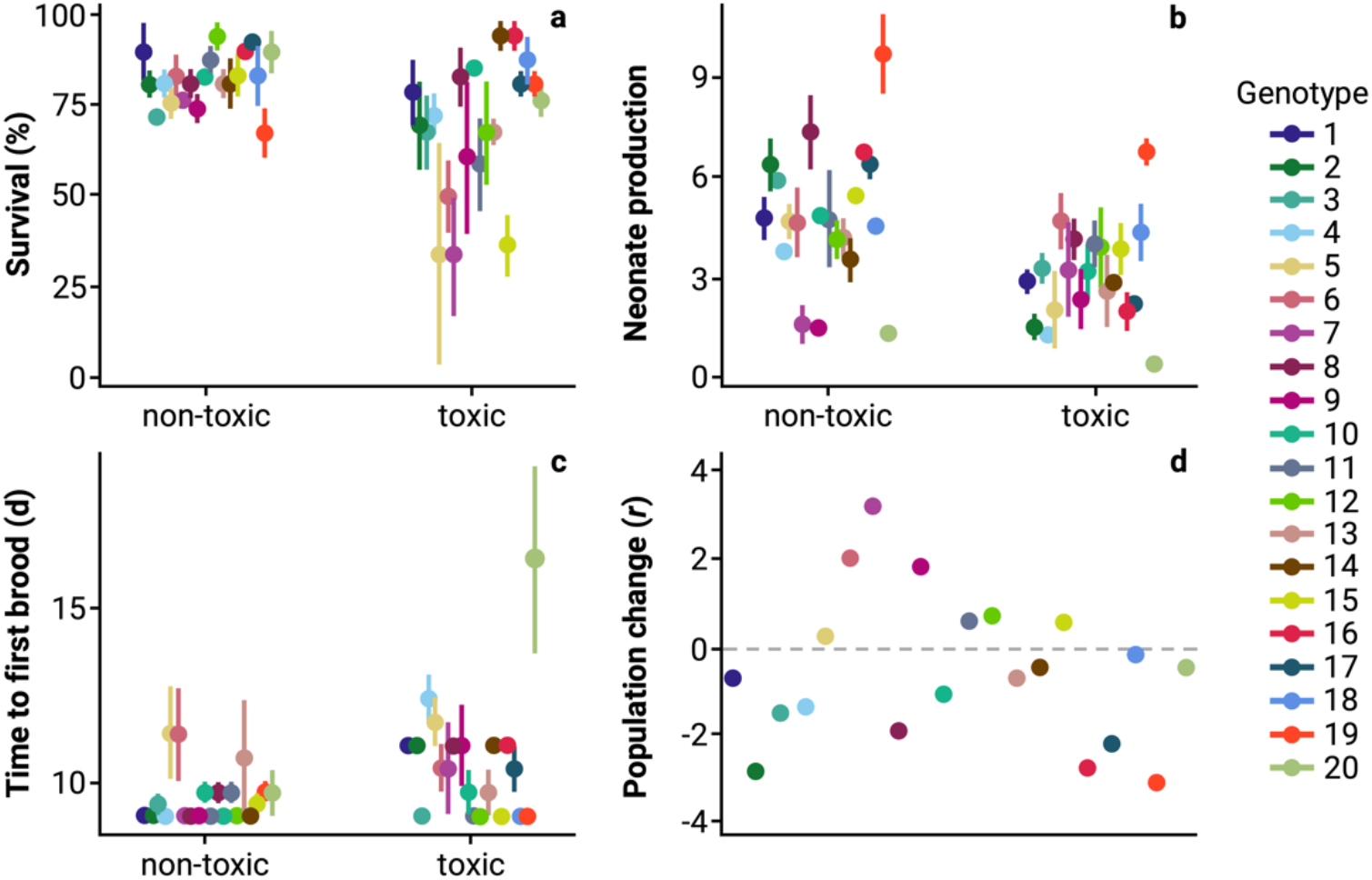
Phenotypic variation among the 20 *Daphnia magna* clones at the end of the 21-d gut microbiome chronic experiment across two treatments: non-toxic diet (Chlorella-only) and toxic diet (3:1 Chlorella:Microcystis). Phenotypes measured were a) survival (%), b) mean number of neonates produced per *D. magna*, and c) mean time to first brood per *D. magna* (d). Panel d) is the exponential rate of population change (*r*) between treatments across each of 20 *D. magna* clonal populations at the end of the 21-d gut microbiome chronic experiment. A positive population growth response is *r* > 0 whereas a negative population growth response is *r* < 0.

To make inferences on the effects of *Microcystis* exposure on population dynamics we constructed Leslie matrices for each clone on both non-toxic and toxic diets. We then calculated the exponential rate of increase (*r*) for each *Daphnia* clonal population on each diet and the mean difference in *r* between toxic and non-toxic for each ‘clone’ and ‘treatment’ combination to determine whether *Microcystis* exposure would have a net positive (*r* > 0), neutral (*r* = 0), or negative (*r* < 0) effect relative to rearing on a control diet (Figure 1d). *Microcystis* exposure negatively impacted 13/20 clones and had positive effects on 7/20 clones (Figure 1d). The mean difference in population growth rate between toxic (2.38) and non-toxic (2.88) for all clones was -0.49, indicating the overall effect of *Microcystis* exposure on *Daphnia* population growth was negative.

### Testing for effects of toxic exposure on Daphnia microbiomes

We found no significant effect of algal treatment (F_38_ = 0.763, P = 0.388) or *Daphnia* clone (F_38_ = 0.265, P = 0.609) on microbiome α-diversity (Figure 2). Effects of algal treatment and clone were likewise modest on ß-diversity; we observed no significant effect of algal treatment (F_1_ = 1.53, P = 0.195) or clone (F_19_ = 1.22, P = 0.173) using Weighted UniFrac (Figure 3a) and a marginally significant effect of clone (F_19_ = 1.17, P = 0.049), but not algal treatment (F_1_ = 0.96, P = 0.505), when assessed using Bray-Curtis (Figure 3b). Using a permutation test for homogeneity of multivariate dispersions, we likewise saw no significant difference in variance in community composition associated with algal treatments (Weighted UniFrac: F_1_ = 0.601, P = 0.443; and Bray-Curtis: F_1_ = 0.009, P = 0.152).

**Figure 2.**
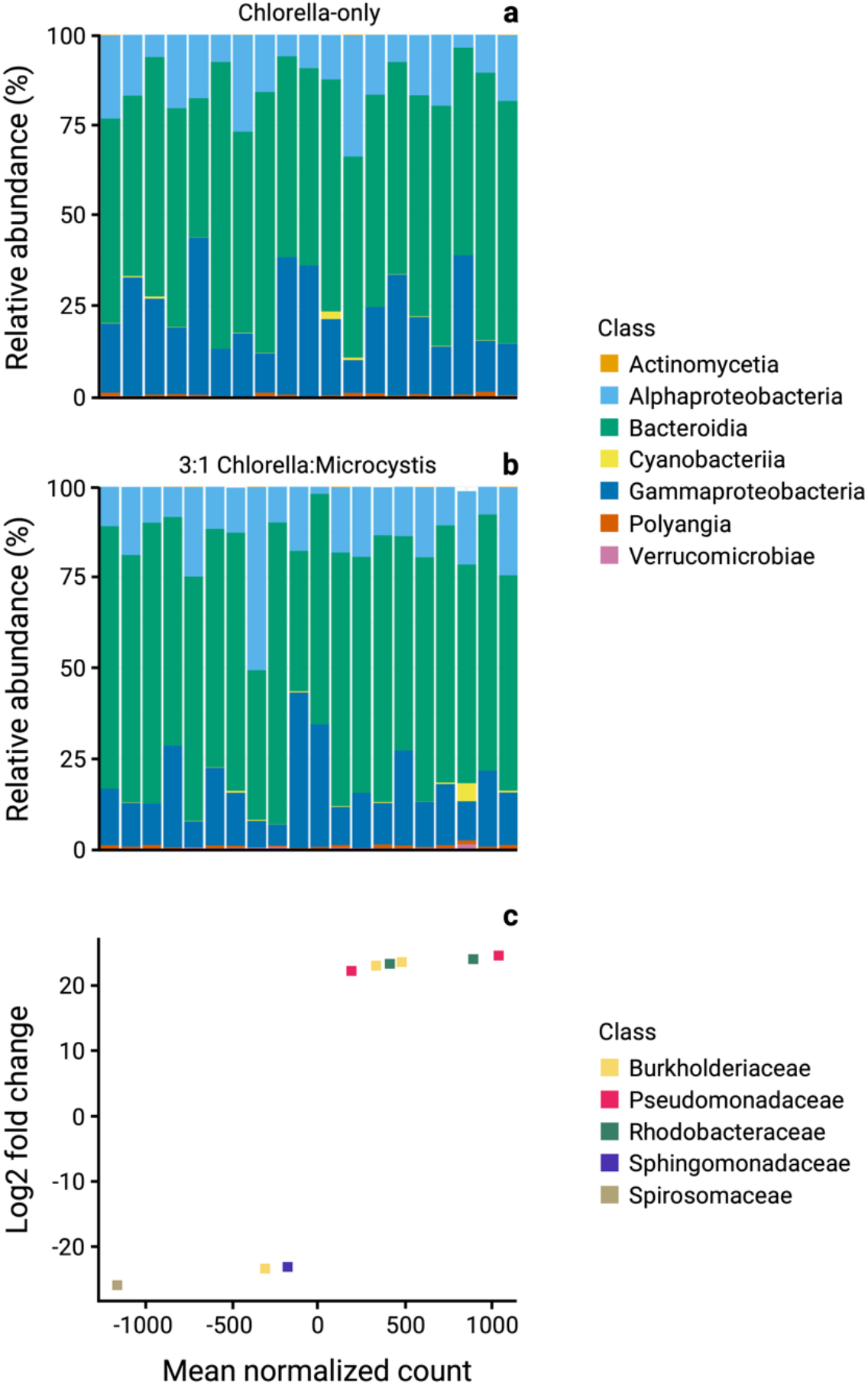
The average relative abundance of bacterial classes in the gut microbiomes of 20 *Daphnia magna* clones across a) non-toxic (Chlorella-only) and b) toxic (3:1 Chlorella:Microcystis) treatments at the end of the 21-d gut microbiome chronic experiment, color coated based on bacterial classes. c) The *DESeq2* differential abundance analysis, color coated based on family.

**Figure 3.**
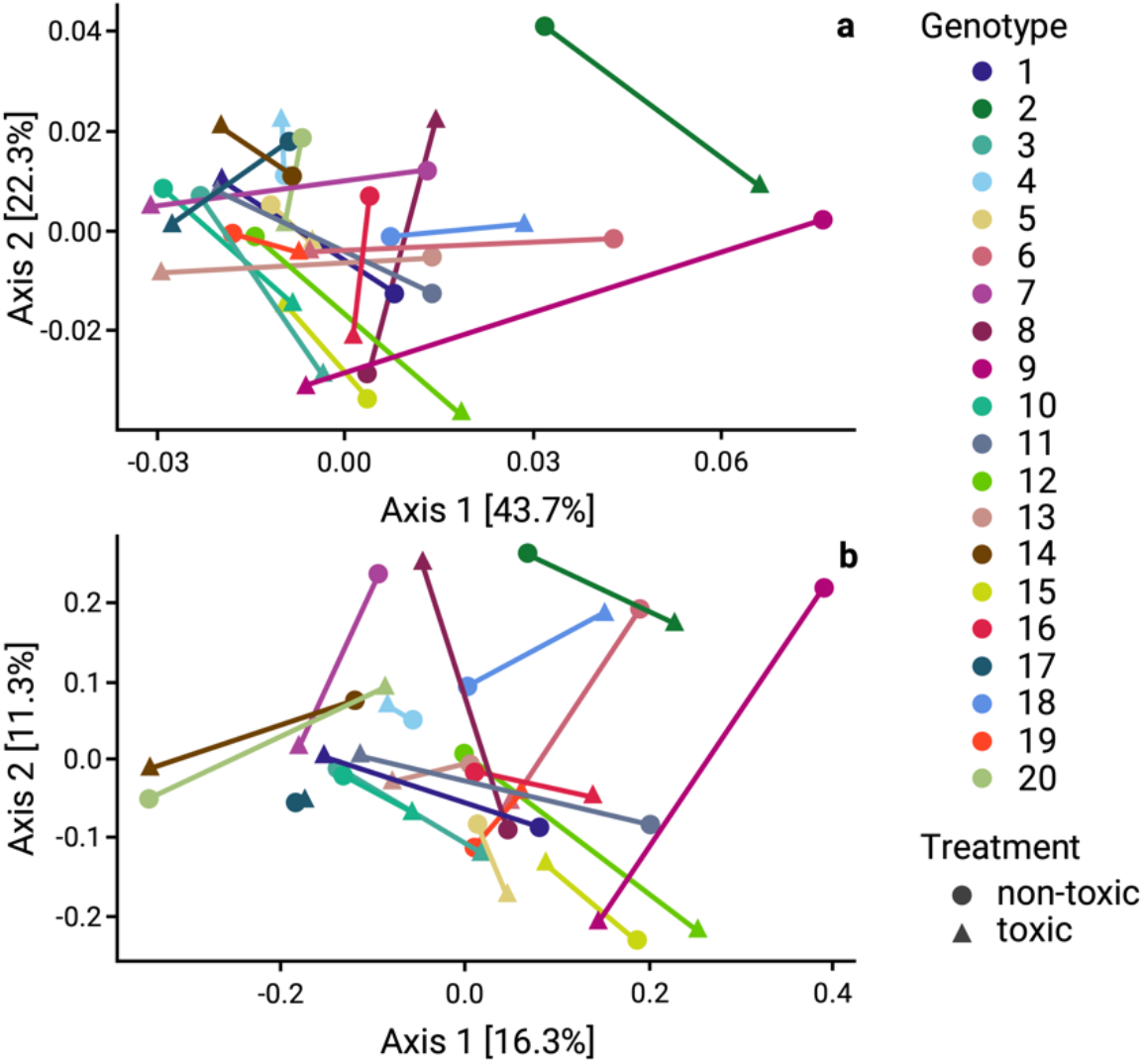
Principal Component of Analysis (PCoA) in a) Weighted UniFrac, and b) Bray-Curtis distances in relation to the gut microbiome across the 20 *Daphnia magna* clones subjected to non-toxic (Chlorella-only; ▲) and toxic (3:1 Chlorella:Microcystis; •) treatments. Each clone is connected by a line to show the shift in host response across treatments.

Although ß-diversity metrics demonstrated no significant community wide differences in microbiome composition, there were particular microbial groups that differed in their abundance across algal diets. A *DESeq2* based analysis of differential read abundance found five differentially abundant bacterial families (Figure 2c): *Spirosomaceae* (log2FC = -25.57, P < 0.001), *Sphingomonadaceae* (log2FC = -23.02, P < 0.001), *Rhodobacteraceae* (log2FC = 23.03, P < 0.001), *Pseudomonadaceae* (log2FC = 23.57, P < 0.001), and *Burkholderiaceae* (log2FC = 23.67, P < 0.001). A *Phylofactor* based analysis identified only *Spirosomaceae* (F_2_ = 11.28, P = 0.001) as a differentially abundant taxa.

To determine whether diet treatments produced parallel shifts in microbiome composition, we employed *multivarvector* (Härer & Rennison, 2022). Treating each clone as a unique contrast, we found the shift in microbiome composition associated with non-toxic and toxic diet exposure was non-parallel (student’s t-test with a null hypothesis of 90° (non-parallel) for both Weighted UniFrac (t_19_ = -0.231, df = 19, P = 0.82, Mean = 89.73) and Bray-Curtis (t_19_ = 0.34, df = 19, P = 0.74, Mean = 90.14) indicated a non-parallel relationship).

### Magnitude of microbial plasticity and degree of parallelism

We next tested whether the magnitude of microbiome shifts that occurred with exposure to toxic diets, as calculated in multivariate space, was associated with the magnitude of host plasticity across diets. To do so, we tested for a correlation between the magnitude of the microbiome shift in multivariate space and the magnitude of change in *r* across diet treatments for each of the 20 clones. We observed no significant relationship between the degree of shifts in the microbiome and the difference in *r* across diet treatments in either Weighted Unifrac (F_18_ = 0.083, P = 0.776, R^2^ = 0.005; Figure 4a) or Bray Curtis (F_18_ = 0.009, P = 0.923, R^2^ = 0.0005; Figure 4b).

**Figure 4.**
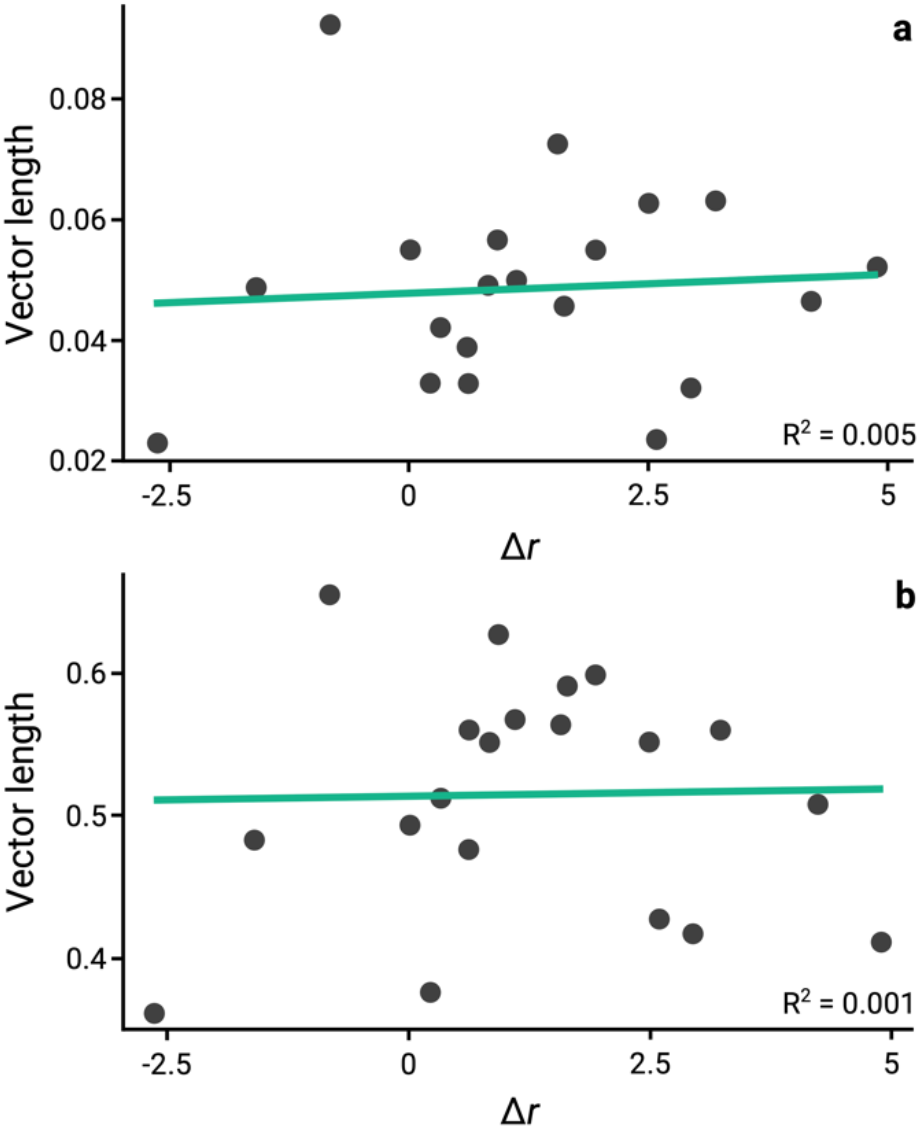
Magnitude of microbial plasticity in a) Weighted UniFrac, and b) Bray-Curtis distances in relation to the difference in difference in population growth rates (*r*) across the 20 *Daphnia magna* clones subjected to non-toxic (Chlorella-only) and toxic (3:1 Chlorella:Microcystis) treatments. The mean vector is represented by the bluegreen line.

## Discussion

### Relationship between phenotypic responses, shifts in the microbiome, and population-level effects

Overall patterns of host phenotypic plasticity associated with *Microcystis* dietary exposure were considerable, with mean shifts in survival, number of offspring produced, and time to first brood generally trending toward a negative response across the 20 *Daphnia* clones (Figure 1a-c). When these phenotypes were combined to parameterize a population projection, dietary exposure to *Microcystis* produced considerable clonal variation and led to an overall decrease in population growth rates (Figure 1d). This response aligns with prior work (Shahmohamadloo et al., 2024), which has shown significant mortality in *Daphnia* due to *Microcystis* (Chislock et al., 2013; Harke et al., 2016; Isanta-Navarro et al., 2021; Shahmohamadloo, Poirier, et al., 2020), along with evidence of adaptive responses to *Microcystis* blooms (Chislock et al., 2013; Hairston et al., 2001; Sarnelle & Wilson, 2005).

In contrast to these pronounced effects on host phenotypic plasticity, *Microcystis* exposure produced only modest shifts in the microbiome (Figure 2a-c), with no parallel changes across *Daphnia* clones (Figure 3). Furthermore, the magnitude of microbiome shifts was not correlated with the magnitude of phenotypic plasticity observed across clones (Figure 4). These findings stand in stark contrast to prior work focused on axenic rearing and microbiome transplants which has suggested the select few community members that do dominate the *Daphnia* gut microbiome (Freese & Schink, 2011)—of which we identified five microbial families that significantly differed (Figure 2c)—have a prominent influence on host fitness including survival, growth, and reproduction when diets feature *Microcystis* (Callens et al., 2018; Macke et al., 2017; Peerakietkhajorn et al., 2016; Sison-Mangus et al., 2015). Our results, however, suggest that exposure to a common environmental stressor did not lead to microbiome-mediated plasticity with adaptive phenotypic or positive demographic effects that could stabilize population growth rates and decrease extinction risk.

### Microbiomes and adaptive plasticity revisited

The role of the microbiome in host responses to environmental stress has been a topic of considerable interest (Henry et al., 2021; Kolodny & Schulenburg, 2020; Rocca et al., 2018; Zilber-Rosenberg & Rosenberg, 2008). Spurred by early work documenting shifts in microbiome composition associated with environmental stress (Engel et al., 2012; Motta et al., 2018), there has been considerable speculation about the role of the microbiome in host acclimation and adaptation (Mueller et al., 2020; Rudman et al., 2019). Our findings raise questions about the circumstances and generality of microbiomes as a mechanism underlying adaptive plasticity in hosts. We show that the *Daphnia* microbiome only exhibits significant shifts in the abundance of five bacterial families. Though it has been proposed that shifts in microbiome composition play a crucial role in influencing the fitness of hosts and stabilizing growth rates to reduce the probability of extinction (Kolodny & Schulenburg, 2020; Mueller et al., 2020; Shade, 2023), we find no correlation between the degree of shift in the microbiome and host adaptive plasticity (Figure 3) or population growth rate (Figure 1d).

Prior work demonstrating a role of adaptive plasticity has largely studied the contribution of microbiomes without host genetic diversity (Baldassarre et al., 2022) or through complete microbiome transplants (Callens et al., 2018, 2020; Houwenhuyse et al., 2021; Macke et al., 2017, 2020). While this work illustrates how influential host-microbiome interactions can be in specific and tightly controlled circumstances, these types of approaches could lead to overestimates of the importance of shifts in the microbiome for host phenotypic responses in more natural settings (Hegg et al., 2021).

An important caveat of our work is that it was conducted under laboratory conditions, where *D. magna* gut bacterial communities can differ significantly from those in natural environments. Recent work shows that microbiome diversity and composition can change rapidly after transfer to the lab, with ongoing shifts even after two years (Houwenhuyse et al., 2023). Despite these differences, key bacterial classes (Hegg et al., 2021; Houwenhuyse et al., 2023)*Burkholderiaceae, Pseudomonadaceae*, and *Sphingomonadaceae* were dominant across field and lab rearing environments and present in high abundances in our study. Our test tests for parallelism (Figure 4) revealed that their relative abundances were not repeatedly changed by exposure to *Microcystis*, aligning with prior work under semi-natural conditions showing no evidence that natural variation in microbiome diversity or composition was associated with tolerance to cyanobacteria (Hegg et al., 2021). Future work that extends our general experimental framework to field conditions is needed to determine whether hosts can modulate microbiome composition through horizontal acquisitions, thereby producing adaptive plasticity and positive effects on population growth.

While it is possible that certain traits or environments might benefit from microbiome-mediate rescue (Mueller et al., 2020; Shade, 2023) or ‘microbiome flexibility’ (Voolstra & Ziegler, 2020), our findings suggest this was not the case here. Moreover, field exposures similarly recover limited evidence for *Daphnia* microbiome shifts in response to *Microcystis*, which does not indicate a strong pathway for adaptive plasticity via microbiome shifts or microbiome rescue (Hegg et al., 2021). Although concepts like microbiome plasticity and rescue were initially motivated by findings in organisms with strong host-microbiome symbiosis, the authors explicitly posited that these phenomena are likely to be broad and widespread (Mueller et al., 2020; Shade, 2023).

In the absence of microbiome rescue or flexibility operating to reduce phenotype-environment mismatches, other mechanisms, such as phenotypic plasticity and adaptation from standing genetic variation, play crucial roles in shaping the phenotypic diversity observed in natural populations (Barrett & Schluter, 2008; Rennison et al., 2019; Rudman et al., 2022; West-Eberhard, 2003). These mechanisms shape the phenotypic response of organisms to environmental challenges (Ghalambor et al., 2007; Price et al., 2003; West-Eberhard, 2003). It is plausible that microbiome-driven adaptive plasticity is also a component of adaptive plasticity but operates on longer timescales (Kolodny & Schulenburg, 2020) or requires transgenerational effects (Baldassarre et al., 2022), as suggested by the microbiota-mediated transgenerational acclimatization concept (Webster & Reusch, 2017). These effects might be limited to systems with high microbiome heritability (Doolittle & Booth, 2017) and may not be as general as initially proposed. Advancing microbiome studies beyond tightly controlled mechanistic investigations to include genetically, environmentally, and phenotypically diverse populations will be critical for understanding the role of microbiomes in organismal responses to environmental change (Greyson-Gaito et al., 2020).

### Quantitative and qualitative replication of host-microbiome studies

Despite observing significant shifts in *Daphnia* life-history traits and population growth associated with *Microcystis* exposure, we observed no community-level differences in gut microbiome composition between *Daphnia* reared on toxic and non-toxic diets. This result aligns with findings from outdoor mesocosms, which demonstrated that despite significant seasonal changes in *D. magna* gut microbiome diversity and composition there was surprisingly no evidence linking natural variation in microbiome diversity or composition with exposure to *Microcystis* (Hegg et al., 2021).

Our results suggest that earlier mechanistic studies—which used axenic cultures and reciprocal microbiome transplants to test the effects of microbiome composition on *Microcystis* toxin tolerance (Houwenhuyse et al., 2021; Macke et al., 2017, 2020)—may not adequately capture the dynamics of host-microbiome interactions in natural settings, and prompts further investigation into the ecological relevance of such mechanistic findings (Greyson-Gaito et al., 2020). These differences highlight major questions about reproducibility and repeatability in microbiome research, due both to underlying stochasticity and inherent differences between experimental environments (Schloss, 2018). In planning this study, we consulted authors on the prior work and they generously shared methodological details which we made efforts to replicate. However, there are experimental differences that could contribute to the considerable variation seen between studies. These include: previous studies did not detail the concentration of *Microcystis* cells or amount of microcystin toxins, and hence exposures may not have been comparable (Orr et al., 2018); biological variation in *Daphnia* test populations and *Microcystis* strains; and inherent differences in lab microbiomes. Whether these differences are sufficient to lead to qualitatively different conclusions about the role of microbiome composition in host fitness is beyond our scope to determine. The differences observed highlight the need to consider not only genetic and environmental factors but also methodological consistency when interpreting microbiome research results. Given the observed differences between studies and the inherent repeatability challenges in microbiome research, our findings should be interpreted within the context of these broader uncertainties. While our results contribute valuable insights, they also underscore the importance of continued investigation and methodological rigor in this field.

### Broader implications

Our findings provide an empirical test of the contribution of the microbiome to host plasticity and population dynamics. Although significant phenotypic changes were observed in *Daphnia* exposed to *Microcystis*, there was no correlation between this response and the degree of change in the composition of the microbiome or any pattern observed indicative of microbiome-mediated rescue. These results suggest that the microbiome’s role in host plasticity may be more context-dependent than previously thought. However, given the scope of this single study, further research is needed to explore how shifts in the microbiome influence reaction norms in fitness associated phenotypes across species and environmental contexts. Future studies in natural conditions, where hosts can acquire a wider range of microbes, will be critical to developing a more comprehensive understanding of how host-microbiome interactions shape natural populations. Such investigations may help resolve longstanding questions about the microbiome’s potential to direct host evolution and serve as a mechanism of rescue in populations facing novel environmental challenges.

## Acknowledgements

We thank S. Porter and the Rudman lab for helpful comments and discussions.

